# Nanopore sequencing from liquid biopsy: analysis of copy number variations from cell-free DNA of lung cancer patients

**DOI:** 10.1101/2020.06.22.165555

**Authors:** Filippo Martignano, Stefania Crucitta, Alessandra Mingrino, Roberto Semeraro, Marzia Del Re, Iacopo Petrini, Alberto Magi, Silvestro G. Conticello

## Abstract

Alterations in the genetic content, such as Copy Number Variations (CNVs) is one of the hallmarks of cancer and their detection is used to recognize tumoral DNA. Analysis of cell-free DNA from plasma is a powerful tool for non-invasive disease monitoring in cancer patients. Here we exploit third generation sequencing (Nanopore) to obtain a CNVs profile of tumoral DNA from plasma, where cancer-related chromosomal alterations are readily identifiable.

Compared to Illumina sequencing -the only available alternative- Nanopore sequencing represents a viable approach to characterize the molecular phenotype, both for its ease of use, costs and rapid turnaround (6 hours).

## MAIN TEXT

Alterations in the copy number of subgenomic regions is one of the characterizing features in many cancers: specific amplifications or deletions can define type and progression of the tumor, and are thus tightly linked to the diagnostic and prognostic process [1–5].

Characterization of cancer genetic features, such as copy number variation (CNV), is typically done on tissue samples, be them surgical resections or bioptic samples; however, the collection of tissue samples is often invasive, harmful and not repeatable [6–10].

On the other hand, liquid biopsy is a non-invasive approach for monitoring tumor features through the analysis of body fluids obtained from cancer patients. The most common liquid biopsy approach for genomic characterization is the analysis of cell-free DNA (cfDNA) from plasma, which can be easily collected at different time-points to closely follow tumor evolution, with limited harm and risks for the patient [6, 11–14]. However, the analysis of cfDNA is challenging as its concentration is very low and cfDNA is highly degraded, with an enrichment of ~169bp fragments. This typical fragmentation pattern is due to the nucleosomes that protect DNA from degradation by DNAses upon its release. Moreover, tumor-derived cfDNA (ctDNA) can be contaminated by healthy DNA due to accidental lysis of blood cell, and its fraction may vary from 0.01% to 60%, depending on tumor features such as tumor volume, stage, vascularization, cell death and proliferation [11, 12].

Third generation sequencing approaches, such as Nanopore technology, interrogate single molecules of DNA and are capable of producing sequences much longer than those generated by second generation sequencing (SGS) methods. The passage of a single DNA filament through a pore produces an electric signal which depends on its sequence; subsequently, the pore becomes available for the sequencing of a new molecule, and the electric signal produced is immediately stored and ready for analysis. This aspect is crucial as it allows the user to obtain sequencing results and perform real-time analyses while the instrument is still running [15, 16].

Unfortunately, as Nanopore technology is optimized for long read sequencing, its protocols are not ideal for analysis of short cfDNA fragments. Moreover, early attempts at sequencing maternal plasma cfDNA for non-invasive prenatal diagnosis resulted in unsatisfactory throughput (< 60k reads) [17]. Thus, before Nanopore-seq potential can be exploited for liquid biopsy applications, effective and standardized workflows need to be developed.

Here we use shallow whole genome sequencing (sWGS) to detect CNVs from cfDNA. sWGS is a read-count based approach that allows detection of genome-wide CNVs from reads produced by a low-coverage (< 1x) whole genome sequencing experiment [18].

To exploit Nanopore technology on cfDNA, we set up a custom protocol through which we sequenced cfDNA from 6 cancer patients and 5 healthy subjects, in both singleplex and multiplex runs (S1, M1 and M2, **Table S1**). Since Nanopore library preparation protocols are designed to enrich long DNA fragments, we have modified the clean-up steps, increasing the ratio of magnetic beads to retain small cfDNA fragments (see supplementary methods).

In this way, we obtained 14,338,633, 19,610,131, and 31,582,051 raw reads from the S1, M1 and M2 runs, respectively: a remarkably higher throughput than previously reported [17] (**Table S1**). Notably, the per-sample throughput was highly variable, even if the amount of input DNA was constant for most of the samples (30ng). Such differences are likely to depend on different efficiencies in the library preparation steps (see supplementary data).

Alignment was performed with both Burrows Wheeler Aligner (BWA) and Minimap2 (see methods). The average percentage of uniquely mapped reads was 98.5% and 85.6%, respectively (**Table S1**). Size distribution of the sequenced cfDNA fragments perfectly matches the fragmentation profile obtained with Agilent Bioanalyzer (**Figure 1**). While Minimap2 is usually recommended for alignment of long Nanopore reads, according to our results BWA is preferable for cfDNA-derived data, probably due to the shorter length of cfDNA fragments.

**Figure 1.**
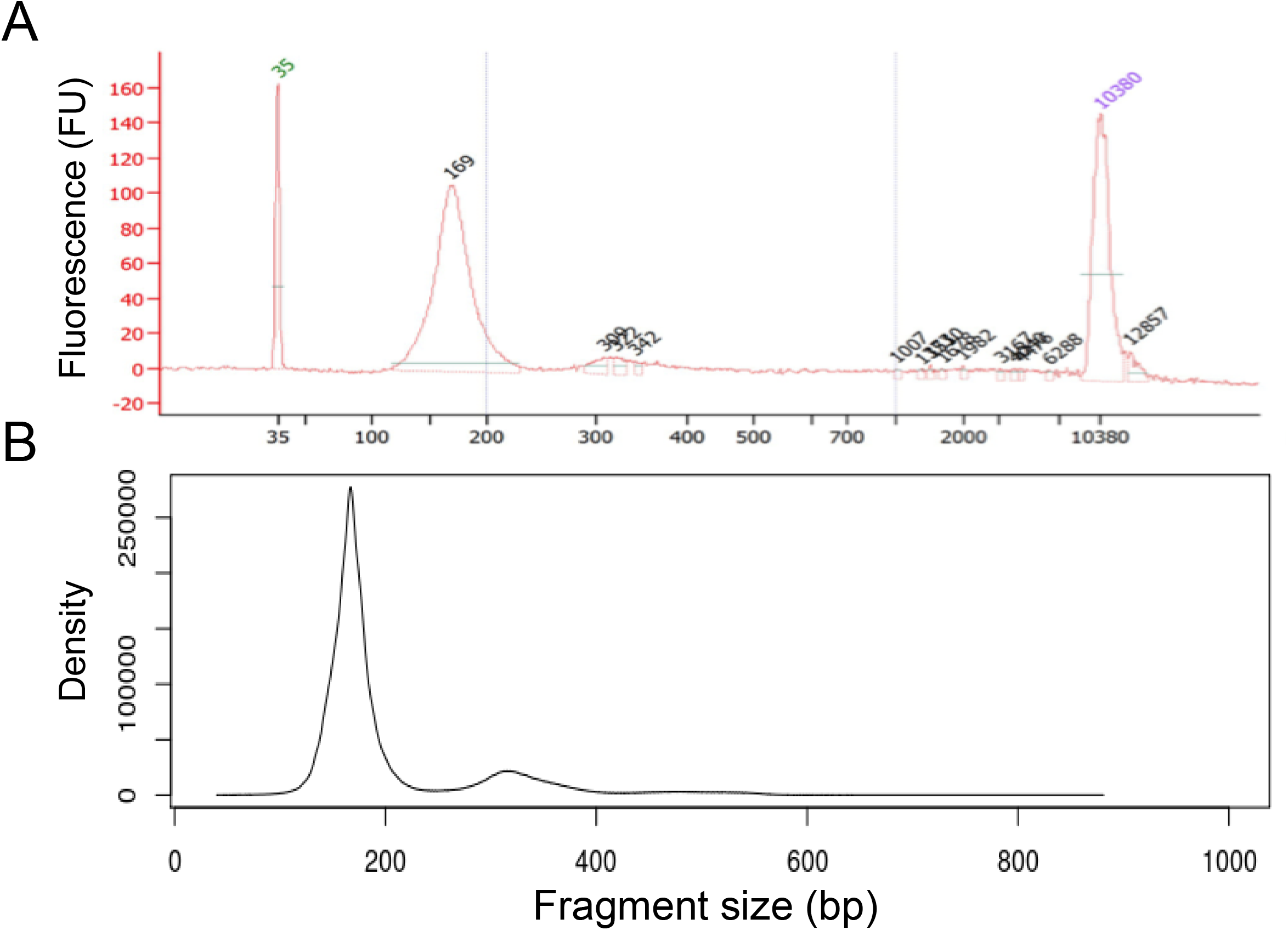
Fragment size distribution. Fragment size distribution estimated through Bioanalyzer (A), and from Nanopore reads (B).

Molecular karyotype of 10 out of 11 samples was successfully produced using NanoGLADIATOR (*“nocontrol”* mode), a recently developed tool to identify CNVs from decrease or increase of read counts (reported as log2ratio) across multiple consecutive windows (bins) [19]. BWA-aligned BAM files were analysed with a bin size of 100k bp, and CNVs have been detected in all the tumoral samples (**Figure 2**).

**Figure 2.**
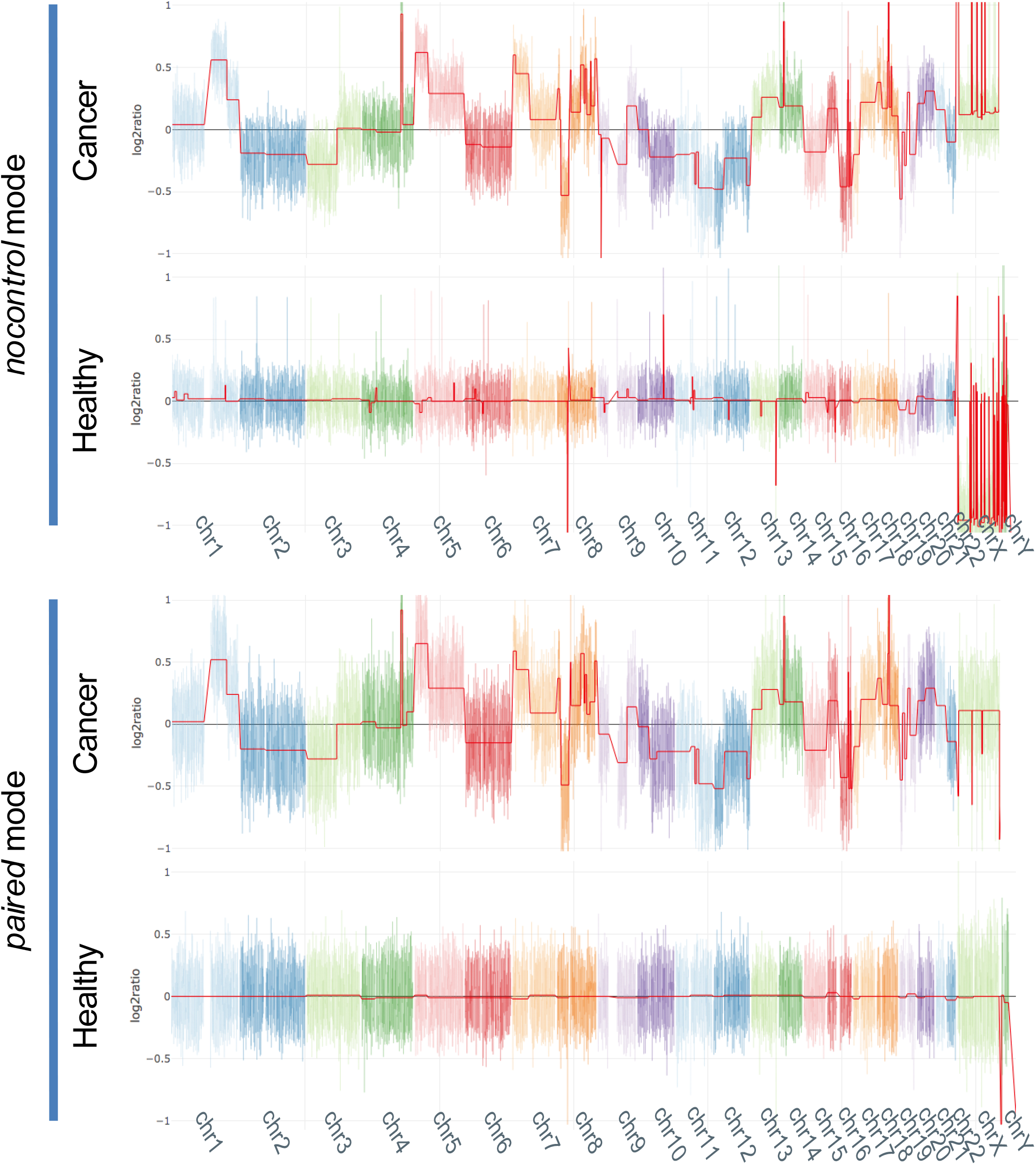
NanoGLADIATOR segmentation plots. Segmentation plots produced with NanoGLADIATOR for samples 19_1231 (cancer) and HM3 (healthy) in *“nocontrol* mode (A), and “*paired”* mode (B). In “paired” mode, HF1 and HM2 were used as controls for respectively l9_l23l and HM3. The red line indicates the segment mean (log2ratio). Each color represents a different chromosome; chromosome Y for sample 19_1231 has been omitted.

Unexpected variations in read-count values were also present in samples from healthy donors (**Figure 2, Figure S1)**. Even though it is possible that these variations represent naturally occurring polymorphisms, this is unlikely: polymorphic variations should present a discrete number of copies (1,3 or 4 copies), which is not the case, as most of these variations have weak log2ratio.

These technical artefacts can be easily filtered out setting a threshold. On the other hand, some of these variations are very similar in terms of length and segment mean (roughly ±0.10) to those we observe in cancer samples and it could be difficult to discriminate real CNVs from these ones (see supplementary data, **Figure 2, Figure S2**). Typically, these artefacts are present in regions containing a higher number of homologous segments, e.g. the sexual chromosomes. Alignment of short reads in such genomic regions is typically challenging and presence of these artefacts is likely due to mapping issues (**Figure S1**) [20]. The fact that most of the variations observed in healthy individuals are shared among samples is another indication that they could represent artefacts (**Figure S1**). In order to minimize the number of artefacts, we used NanoGLADIATOR in *“paired”* mode, which generates segmentation results comparing test samples with a control sample. In “paired” mode, we used as control a merged BAM files from the samples of healthy donors (see supplementary methods) that allowed us to decrease the log2ratio of these artefacts to ±0.04 and, consequently, to drastically increase the specificity of the analysis (**Figure 2, Table S2**). We then compared the performance of Nanopore sequencing with a standard SGS approach by analysing four of the tumoral samples through Illumina sequencing (17-24M, 150bp single end reads, see methods).

Illumina and Nanopore results (*“nocontrol”* mode) were strongly correlated (R = 0.93 – 0.99, *p* << 0.001), with concordant log2ratio values at 95-98% of the genomic positions (**Figure 3, Table S3**). To assess the performances of our approach at even lower sequencing depth, we subsampled the BAMs to 2M raw reads: the results obtained are highly concordant with the full-depth BAMs (R = 0.93 – 0.99, *p* << 0.001, 94-99% concordant bins, **Figure 3, Table S4**).

The marginal loss of performance observed is comparable to the one obtained when subsampling Illumina data (**Table S5**).

**Figure 3.**
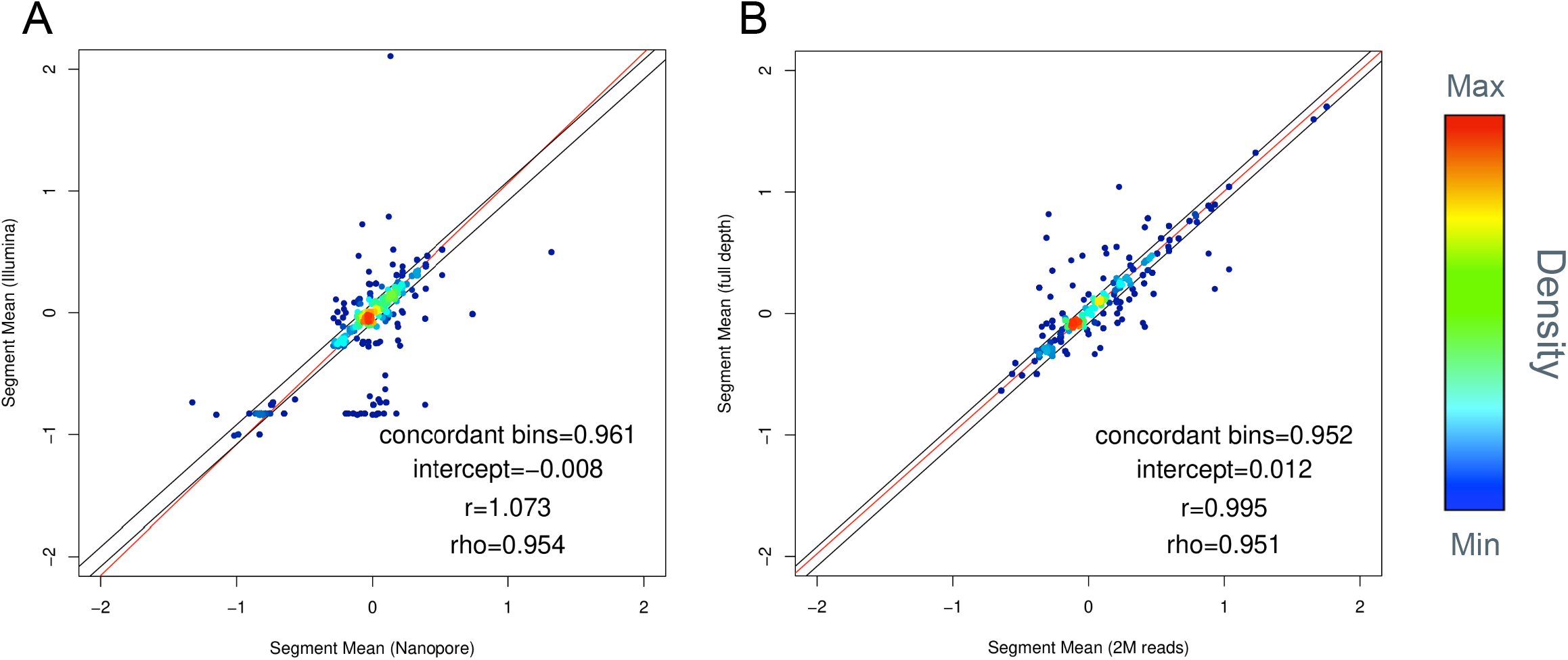
Comparison of segmentation results. (A) Correlation of Nanopore and Illumina segment mean values (sample 19_744); (B) Comparison of Nanopore segment mean values from the full-depth BAM file and from a 2M reads subsampled dataset (sample 18_1130). Each genomic bin is represented as a dot, colours indicate dot density. Regression lines are shown in red. Black lines indicate the thresholds for concordant bins.

Since the ultimate aim of the analysis is to obtain information on the tumour, we next assessed the status of genes commonly altered in lung cancer (**Figure 4**) [21–27]. Indeed, pathogenetic CNVs were readily observed, with EGFR amplification prominently present in all samples, and most of other genes altered in at least two samples. Many of these structural alterations directly affect progression of the cancer and therapeutic options. For example, RICTOR amplification identifies a subgroup of lung cancer and its presence has been linked to the response to mTOR inhibitors [25]. Similarly, MYC amplification confers resistance to pictilisib in models and PIK3CA amplification is associated with resistance to PI3K inhibition [28, 29] in mammary tumors.

**Figure 4.**
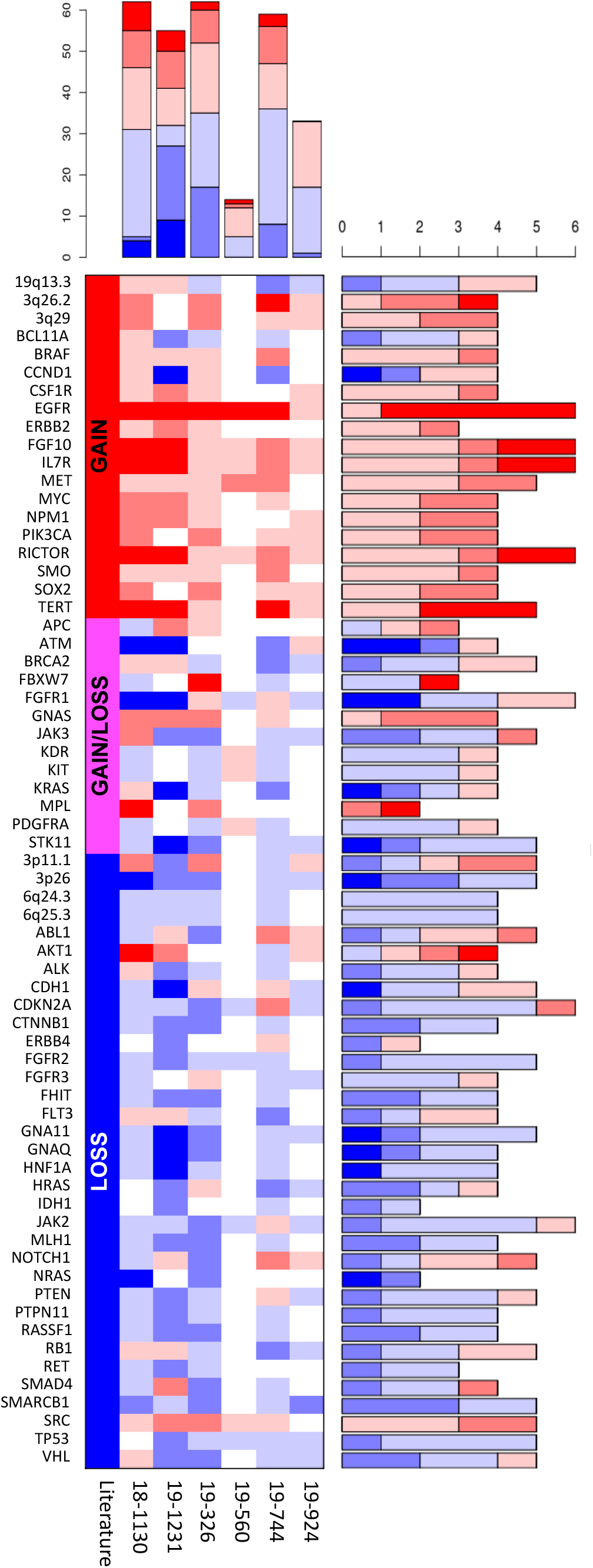
Landscape of clinically-relevant copy number variants. Copy number variants of specific genes (rows) are shown for the individual patients (columns). The shading indicates levels of amplification (red tones, 0.04-0.15, 0.15-0.30, >0.30 log2ratio) and deletion (blue tones, 0.04-0.15, 0.15-0.30, >0.30 negative log2ratio). The top and right barplots show the number of CNVs in one patient and the number of patients with CNVs for a given gene, respectively. The expected status for a given gene based on the literature is shown in the left side.

Our report is the first successful attempt to obtain a CNV profile from plasma cell-free DNA of cancer patients using Nanopore technology. Our results show that Nanopore sequencing has the same performance of SGS approaches and, in terms of throughput and sequencing costs, it is comparable to an Illumina MiSeq (V3 reagents, 22-25M single-end reads).

MinION is the entry-level sequencer by Nanopore technology, and its cost is extremely low (~1000 euros) compared to SGS sequencers whose price is in the order of tens of thousands of euros. Reduced overall instrumentation costs makes this approach accessible to most of the research groups which would otherwise be forced to outsource the sequencing, or to gain access to shared sequencers, leading often to long queues and delays. Moreover, SGS is cost effective only when dealing with a large number of patients. This aspect is crucial with regards to clinical analyses, as it leads to a centralization of sequencing-based assays, which are mainly performed in big hospitals that collect samples from large geographic areas.

On the contrary, Nanopore technology is extremely scalable, and only a modest number of patients is required in a multiplexed run, leading to short recruitment times and, consequently, faster results. As we demonstrate that reliable results can be obtained from as few as 2M reads. Based on the throughput obtained in our study, it should be possible to analyse up to 7-15 patients in a single run. Since reads are stored as soon as they are produced, they can be analysed while the experiment is still running by taking advantage of the real-time mode of NanoGLADIATOR.

This feature might come useful when analysing single samples, especially in those patients with lower fraction of ctDNA, for which a higher number or reads and, consequently, a higher resolution may be preferable. In such a context, it would be possible to inspect the CNV profile while the run is still ongoing, and stop once the desired resolution is reached, saving the sequencing power of the flow cell, which can be washed and reused for other samples.

According to our sequencing statistics, 2M reads are produced in less than 3 hours. This means that the entire workflow-from blood withdrawal to bioinformatic analyses-can be performed in less than a working day. This is something unique to Nanopore sequencing, as SGS approaches based on sequence-by-synthesis technologies make reads available only at the end of the whole run, which can last days.

All these features represent advantages over current sequencing technologies and might drive the adoption of molecular karyotyping from liquid biopsies as a tool for cancer monitoring in clinical settings.

## MATERIALS AND METHODS

### Sample collection and cfDNA isolation

Blood from 5 unrelated healthy donors and 6 unrelated metastatic Non Small Cell Lung Cancer patients was collected in EDTA vacuum tubes. Blood samples were centrifuged at 1600g x 10”, and plasma was carefully collected with a pipet without disturbing sedimented blood cells. cfDNA was extracted from 4ml of plasma using QIAamp Circulating Nucleic Acid Kit (QIAGEN, 55114), it was quantified via Qubit Fluorometer (Thermo Fisher Scientific, dsDNA HS assay kit, Q32851), and its fragmentation pattern was obtained via Agilent 2100 Bioanalyzer (Agilent, High Sensitivity DNA kit, 5067-4626). Extracted cfDNA was stored at −80° C.

### Nanopore library preparation and analysis

For library preparation EXP-NBD104 and SQK-LSK109 protocols were used: the bead/sample ratio of AMPure XP beads (Beckman Coulter, A63880) was increased to 1.8x in all clean-up steps. All the other steps were performed following the manufacturer’s instructions.

The SQK-LSK109 protocol was used for the run S1. In the case of the multiplex runs M1 and M2, 25ul of each barcoded sample were pooled together before adapter ligation. The pool was then cleaned-up using 2.5X AMPure XP beads.

S1, M1 and M2 runs were performed using FLO-MIN106 (R9.4) flow cells on a GridION sequencer. FASTQ files were generated via real-time high-fidelty basecalling during the run, and Porechop (https://github.com/rrwick/Porechop) was used to de-multiplex FASTQ files of multiplex runs (M1, M2), and to trim adapters of all the runs.

Minimap2 (with -*ax map-ont* flags) [30] and BWA mem (with -*x ont2d* flags) [31] were used to align raw reads, using the human_g1k_v37_decoy as reference genome.

The CIGAR field of aligned BAMs was used to determine fragment length of sequenced cfDNA (**Figure 1**).

NanoGLADIATOR was used to generate molecular karyotypes of BWA aligned BAMs with a bin size of 100kb [19].

For “paired” mode analysis, HF1 was used as a control for female patients, and BAMs from HM2 and HM3 were merged and used as control (Healthy_Males_Pool, HMP) for male patients (see supplementary information).

Additional details on patients features, library preparation and run statistics are summarized in **Table S1**.

### Illumina library preparation and analysis

Illumina libraries for samples 19_924, 19_744, 19_1231 and 18_1130 were prepared from 15ng of input DNA, using Ovation Ultralow V2 DNA-seq Library Preparation Kit (NUGEN, 0344NB-A01), sequencing runs (150bp, paired end) were performed on a NovaSeq 6000 sequencer (Illumina).

Only R1 reads were used for CNV analysis treating them as the product of a single end sequencing experiment, in order to simplify subsequent steps such as subsampling and comparison with Nanopore results. This strategy doesn’t introduce any methodological bias, since Illumina singleend and paired-end CNV results are highly correlated (**Table S5**).

FASTQ files were aligned with BWA mem using human_g1k_v37_decoy as reference genome.

XCAVATOR was used to generate molecular karyotypes of BWA aligned BAMs with a bin size of 100kb [32].

### Segmentation comparison

Custom R scripts were used to compare segmentation results:

When comparing two experiments, the “segment mean” value of each of the 100kb bins was correlated (corr.test function, R base package, *method=”spearman”*).

To determine the percentage of genomic positions with concordant copy number status, we considered two bins as “concordant” if their segment mean differs by ±0.08.

Chromosome Y bins were ignored when analysing female patients.

When comparing Illumina and Nanopore results, even if the bin size used was the same, the starting positions of the bins slightly differs among the two pipelines; an XCAVATOR bin is considered corresponding to a NanoGLADIATOR bin if its starting position falls between the starting and the end position of the NanoGLADIATOR bin.

Only NanoGLADIATOR bins for which it was possible to identify a corresponding XCAVATOR bin were considered for subsequent analysis.

## Supporting information

Fgure S2

## Author contributions

Conceptualization: FM, SGC; patient recruitment and DNA isolation: IP, MDR, SC; sequencing experiments: FM, AM; formal analysis, investigation, and software: FM, AM, RS; visualization: FM; writing: FM, SGC.

## Data and materials availability

Sequencing data are available upon request.

---

**Figure S1. Technical artefacts in healthy samples**

(A) Shared CNVs in healthy samples HF1 and HM3 are indicated by blue squares. (B) Venn diagram reporting recurring genomic bins with altered log2ratio in healthy samples.

**Figure S2. Segment mean and segment length of Nanopore results**

Correlation of segment mean and length in nocontrol (A) and paired mode (B). Every dot represents a segment. Segment mean is reported on the x-axis and segment length (number of bins per segment) on the y axis. Vertical lines indicate the threshold used to discriminate artefacts from CNVs (log ratio ±0.04). The lower range of the segments is shown in the lower plot for each sample.

**Table S1. Case series and run statistics**

**Table S2. Performance of NanoGLADIATOR pipeline in “nocontrol” and “paired” mode**

**Table S3. Correlation of Illumina and Nanopore results**

**Table S4. Correlation of Nanopore results: subsampled BAMs (2M reads) Vs full depth BAMs (“nocontrol” and “paired” mode)**

**Table S5. Correlation of Illumina results: paired-end Vs single-end, and subsampled BAMs (2M reads) Vs full depth BAMs.**

## SUPPLEMENTARY INFORMATION

### Throughput variability

To assess the effects of input DNA on per-sample throughput, we performed library preparation of samples HM2, HM1 and HF1 with respectively 15, 30 and 60 ng of DNA; however, the amount of reads produced was very consistent among the three samples (~3M reads, **Table S1**), suggesting that input DNA has a low impact on the final throughput.

For the run M2, we quantified eluted DNA after each clean-up step via Qubit Fluorometer: Since DNA concentration highly correlates with read yield, differences in per-sample yields are likely attributable to a different efficiency of library preparation steps rather than amount of input DNA. Nanopore protocols suggest pooling equimolar quantities of barcoded samples prior to adapter ligation to avoid differences in per-sample throughput. However, in order to avoid any waste of DNA and aiming at obtaining the maximum amount of reads from a single flow-cell, we loaded the entire barcoded sample for each patient, which may explain the observed variability. Unexpectedly, the relative-throughput (sample reads/total run reads) of cancer patients is remarkably higher compared to healthy subjects (**Table S1**).

Since there were no differences in input DNA, and per-sample throughput depends mainly on library preparation efficiency, it is possible that the presence of ctDNA positively affects library preparation efficiency; however, the biological aspects of this behaviour are not clear and should be further investigated.

### Artefact filtering using NanoGLADIATOR in *“paired”* mode

False positives observed in healthy subjects are similar to cancer patients’ CNVs in terms of length and segment mean; hence, it would be challenging to set up filtering criteria to discriminate them from real positives (**Figure S2**).

Most of the variations observed in healthy donors are shared by at least 2 healthy subjects, suggesting that they may be errors introduced by the technique itself rather than patient-specific alterations (**Figure S1**).

We used NanoGLADIATOR in *“paired”* mode with the aim of correcting eventual method-specific artefacts: we tested it on healthy male subjects, using each sample as both case and control, in any possible combination.

Using this strategy, we were able to remove 82-100% of false positive bins in healthy males samples (segment mean threshold ≥ 0.04, or <=−0.04) (**Table S2, Figure S2**).

Notably, when HM1 was not used neither as a case nor as control, the number of false positives is reduced by 100% (**Table S2**), suggesting that it may be enriched in sample-specific artefacts; hence, it has not been used as a control in subsequent *“paired”* analyses to avoid the introduction of biases.

HM2 and HM3 BAM files have been merged and the resulting BAM (HMP) has been used as control for male patients, while HF1 has been used as control for female patients.

This approach doesn’t negatively affect the performance of the analysis, as the number of copynumber altered bins is reduced by less than 5% in most of the tumoral samples and increases by 29% in sample 19_744; sample 19_560 is the only exception, with a reduction of roughly ~40% (**Table S2**).

19_560 shows the lowest number of altered bins and the lowest segment mean standard deviation (calculated on autosomes) (**Table S2**).

A lack of clonal CNVs in the tumor, or a lower concentration of ctDNA fragments among the overall cfDNA population can explain these observations; it is therefore not surprising to observe an “healthy-like” genotype, with false positives representing a large part of the detected CNVs.

