## Supplementary material for "Nanopore sequencing from liquid biopsy: analysis of copy number variations from cell-free DNA of lung cancer patients": Figure S1

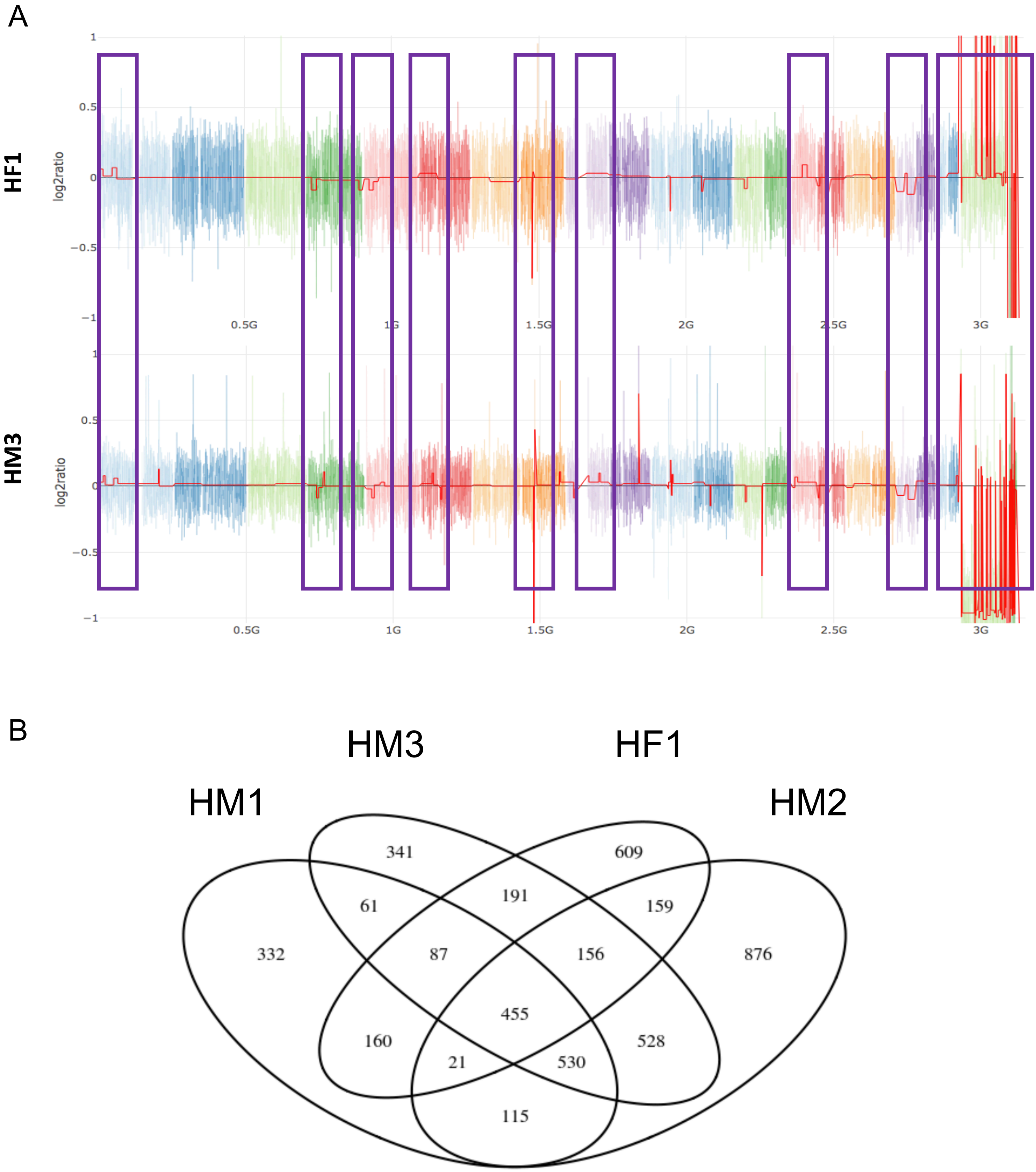


**Figure S1. Technical artefacts in healthy samples**
(A) Shared CNVs in healthy samples HF1 and HM3 are indicated by blue squares. (B) Venn diagram reporting recurring genomic bins with altered log2ratio in healthy samples.
