## Supplementary material for "Nanopore sequencing from liquid biopsy: analysis of copy number variations from cell-free DNA of lung cancer patients": Fgure S2

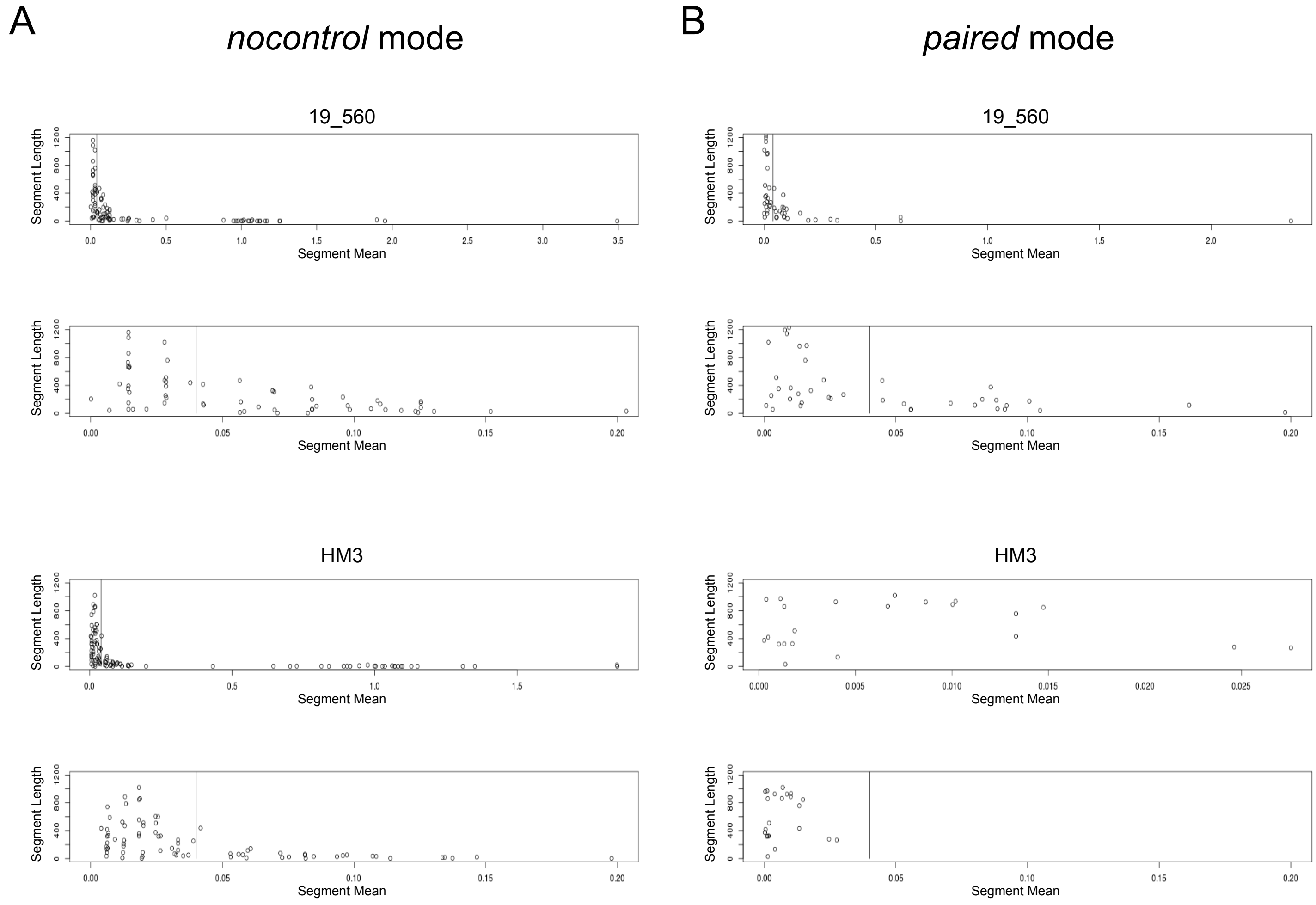


**Figure S2. Segment mean and segment length of Nanopore results**
Correlation of segment mean and length in nocontrol (A) and paired mode (B). Every dot represents a segment. Segment mean is reported on the x-axis and segment length (number of bins per segment) on the y axis. Vertical lines indicate the threshold used to discriminate artefacts from CNVs (log ratio ±0.04). The lower range of the segments is shown in the lower plot for each sample.
